# Threshold-like transcriptomic responses of European beech to simulated future climates

**DOI:** 10.64898/2025.12.16.694614

**Authors:** P Wolf, T Bhatia, BG Jákli, M Baumgarten, C Lindermayr, F Johannes

## Abstract

Beech *(Fagus sylvatica)* is one of the most important forest tree species in Europe, both in terms of land coverage and ecosystem services. Although it is projected to be strongly affected by climate change, insights into its functional genomic responses to realistic future scenarios are still lacking. We conducted a fully controlled ecotron experiment to examine the transcriptomic responses of 125 young, naturally regenerated beech trees exposed to regionalized dynamic climate series representing a reference period (1987–2016) and two future climate scenarios (Representative Concentration Pathways, RCP2.6 and RCP8.5, for 2071–2100) in Germany. RCPs describe alternative greenhouse gas trajectories established by the IPCC, ranging from strong mitigation (RCP2.6) to high, unmitigated emissions (RCP8.5), the latter representing a worst-case future scenario. Although both scenarios produced substantial transcriptional changes relative to the reference period, RCP8.5 elicited markedly stronger responses than RCP2.6, with nearly twice as many differentially expressed genes and unique transcriptional programs across functional categories that were not predictable from the RCP2.6 transcriptomes. Our study reveals emergent functional responses in European beech under projected late 21st-century climate conditions in Germany and highlights potential biomarkers for monitoring ecosystem responses to climate change.

---

Human activities have raised global temperatures by ∼1.1 °C since pre-industrial times, with current policies projecting ∼2.7 °C warming by 2100 (IPCC 2023; UNEP 2023). Atmospheric CO_2_ already exceeds 410 ppm and may surpass 1000 ppm by 2100, while tropospheric ozone (O_3_) in Europe has tripled since pre-industrial times (IPCC 2023; Karlsson et al. 2017; Cooper et al. 2014). Since future atmospheric composition will depend on socio-economic development, land use, and policy, Representative Concentration Pathways (RCPs) were developed to capture possible trajectories, ranging from strong mitigation (RCP2.6) to unabated emissions (RCP8.5) (van Vuuren et al. 2011). The primary effects on plants of elevated CO_2_ have been extensively studied, and show increased photosynthetic rates, growth, partial stomatal closure and improved water-use-efficiency (van der Kooi et al. 2016; Cernusak et al. 2019; Ainsworth und Long 2005). O_3_ is both a greenhouse gas and a phytotoxic pollutant that enters leaves via stomata and decomposes in the apoplast, generating reactive oxygen species (ROS). Excess ROS overwhelm antioxidant defences, leading to oxidative stress, cellular damage, reduced photosynthesis and growth (Vollenweider et al. 2003; Günthardt-Goerg und Vollenweider 2007; Gupta et al. 2015).

Forests are highly vulnerable to rapid climate change, with evidence of declining function (IPCC 2023; Seidl et al. 2017). For European beech (*F. sylvatica*), simulations project 20–50% growth losses under RCP8.5 across much of its range, with major impacts on ecosystem services and carbon storage (Martinez Del Castillo et al. 2022). Given its ecological and economic importance, *F. sylvatica* has been widely studied under elevated O_3_ (Olbrich et al. 2005; Betz et al. 2009), drought (Müller et al. 2017; Cuervo-Alarcon et al. 2021), and CO_2_ enrichment - (Košvancová et al. 2009; Paffetti et al. 2011) with FACE experiments providing key insights under semi-natural conditions (Olbrich et al. 2010; Han et al. 2011). However, most studies have generally focused on a narrow set of physiological traits or limited gene targets and with the only recent availability of a reference genome (Mishra et al. 2018), transcriptomic investigations remain rare (Garosi et al. 2022).

Because plants respond to multiple interacting drivers, single-factor studies may miss emergent responses. Controlled environment facilities (CEFs) address this by simulating realistic multivariate conditions (Boeck et al. 2015; Hanson und Walker 2020; Ogle et al. 2021) and are therefore critical for understanding adaptation under future scenarios. Robust simulations require high-resolution climate inputs; however, while global models (GCMs) project large-scale trends, their coarse resolution (100–250 km) limits regional use. Dynamically downscaled regional models (RCMs), refined with local observations, provide the temporal and spatial detail needed for accurate impact assessments (Vanderkelen et al. 2020; Clobert et al. 2018; Jákli et al. 2021b).

To investigate how *F. sylvatica* responds at the molecular level to projected climatic changes, we conducted a controlled-environment experiment using dynamically downscaled regional climate simulations representative of a Central European forest site (German Spessart). Young trees derived from local natural regeneration were grown for two consecutive seasons under precisely controlled conditions in TUMmesa ecotron chambers (Jákli et al. 2021b; Roy et al. 2021), where hourly meteorological inputs for temperature (t_a_), relative humidity (rh), global radiation (R_g_), air pressure (P), ground-level ozone (O_3_), and carbon dioxide (CO_2_) were used to reproduce present and future climatic conditions with realistic seasonal dynamics (Fig. 1a). Four climate regimes were applied: a present-day control (PC; O_3_ = 39 ppb, CO_2_ = 370 ppm, 14.6 °C, an ozone-enriched present climate (PC_High; O_3_ = 44 ppb, CO_2_ = 370 ppm, 14.6 °C), and two future climate scenarios RCP2.6 (O_3_ = 42 ppb, CO_2_ = 416 ppm, 15.5 °C) and RCP8.5 (O_3_ = 49 ppb, CO_2_ = 924 ppm, 17.7 °C) representing the period 2071–2100 (Fig. 1a). Leaf transcriptomes were analyzed after two growing seasons to identify genes and pathways responsive to these contrasting climates (Methods). By integrating differential expression and network analyses, we aimed to determine whether transcriptomic responses scale predictably with climate severity or whether extreme conditions elicit qualitatively distinct molecular programs.

**Figure 1:**
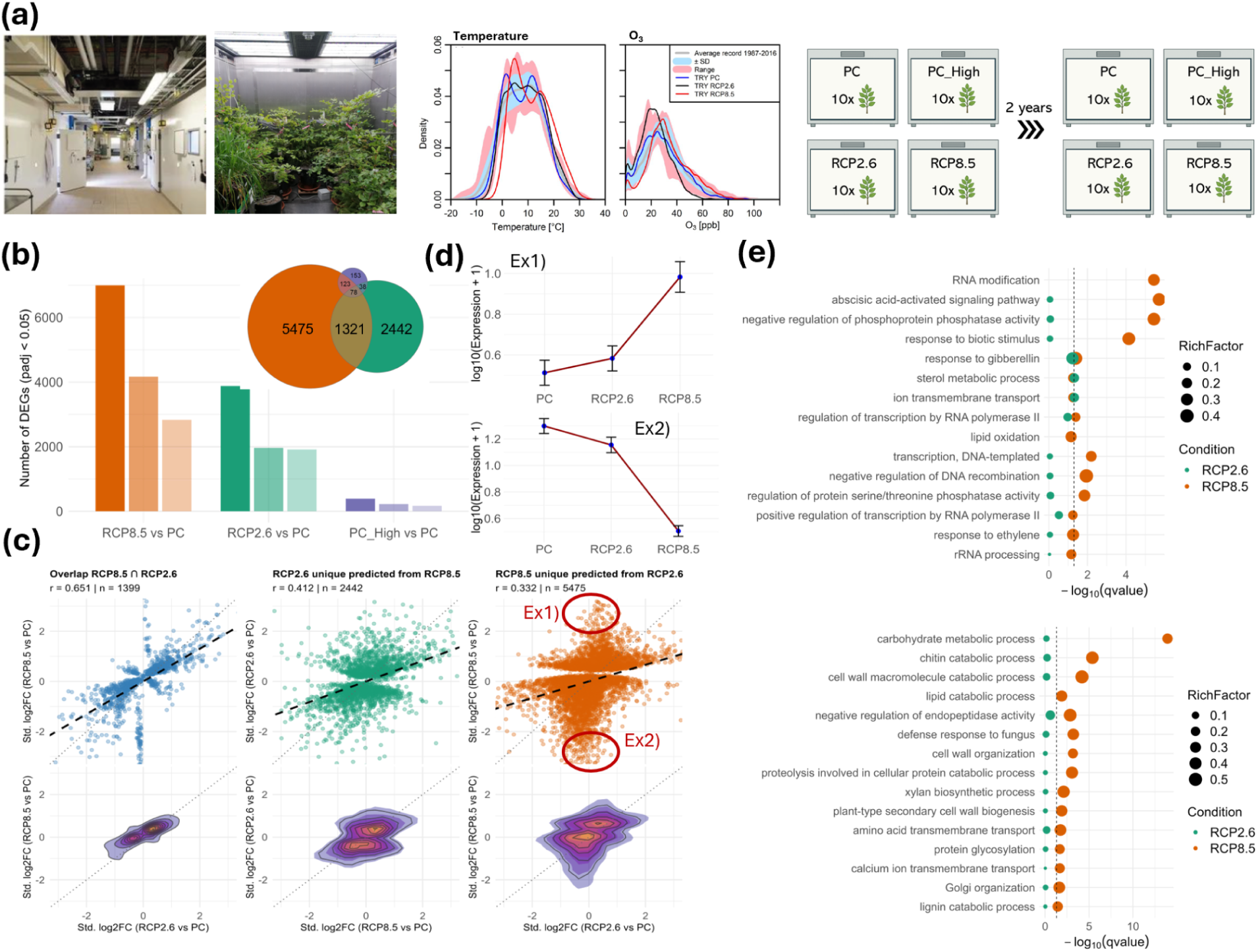
Evidence for threshold-like transcriptional responses. **(a)** Left panel:TUMmesa-ecotron walk-in climate chambers. Middle panel: Probability density functions for the Test Reference Years (TRY) and the mean annual cycles for temperature and O_3_ recorded at the climate station „Spessart“ between 1987-2016. Historically observed ranges are shown in red. Source: (Jákli et al. 2021b). Right panel: Experimental setup - Ten young Fagus sylvatica trees from natural regeneration were grown in fully controlled ecotron environments for two years under a present climate (PC, O_3_=39ppb, CO_2_=370 ppm, 1987–2016), present climate with elevated ozone (PC-High, O_3_=44 ppb, CO_2_=370 ppm), a moderate (RCP2.6, O_3_ =42 ppb, CO_2_ =416 ppm) and a drastic future climate scenario (RCP8.5, O_3_ =49 ppb, CO_2_ =924 ppm). **(b)** Total number (left) of significant (padj < 0.05) DEGs, upregulated (LFC > 0, right) and downregulated (LFC < 0, middle) DEGs. Venn visualization of the overlap of DEGs. **(c)** Linear correlation and density plot of standardized (z-transformed) expression fold-changes compared between scenarios. Genes were considered unique if they were identified as DEGs exclusively within the given scenario. **(d)** Mean expression values from all genes between Ex1) −1 < Standardized log_2_FC (K4 vs K7) < 1 and Standardized log_2_FC (K2 vs K7) > 2; and Ex2) −1 < Standardized log_2_FC (K4 vs K7) < 1 and Standardized log_2_FC (K2 vs K7) < -2. **(e)** GO (BP) enrichments for up- and downregulated genes in RCP8.5 (padj < 0.05, LFC > 0, LFC <0, respectively) and their corresponding enrichment in RCP2.6.

## Non-linear transcriptomic shifts of *Fagus sylvatica* under severe future climate

Across the four simulated climate regimes, *F. sylvatica* showed strong differences in leaf transcriptomic activity. Both the number and identity of differentially expressed genes (DEGs) changed sharply under future conditions. We detected 3,879 DEGs (1,914 up-and 1,965 downregulated) under the moderate RCP2.6 scenario and 6,997 (2,829 up-and 4,168 downregulated) under the severe RCP8.5 scenario, each relative to the present-climate control (PC-PC; *padj* < 0.05; Fig. 1b). The ozone-enriched present climate (PC-High) yielded only 392 DEGs (168 up- and 224 downregulated), showing that elevated O_3_ alone caused relatively minor transcriptional changes.

Examining the DEG overlaps between conditions revealed both shared and scenario-specific responses (Fig. 1b). A total of 1,321 DEGs occurred in both RCP2.6 and RCP8.5, and likely represent a core set of genes involved in general acclimation to future climates. However, more than 5,000 DEGs were unique to RCP8.5, suggesting activation of additional regulatory and stress-related pathways under severe conditions. Overlap with the ozone-only treatment was minimal (<5%), which indicates that the broad transcriptomic shifts under the RCP scenarios were largely distinct from those induced by ozone alone.

To test whether transcriptional responses are scaled with climatic severity, we compared standardized expression fold-changes between scenarios. We found that expression shifts in DEGs unique to RCP8.5 were poorly predicted from those in RCP2.6 (*r* = 0.332, *n* = 5,475; Fig. 1c-d), suggesting that severe responses were not a linear amplification of moderate ones. The reverse comparison produced a stronger correlation (*r* = 0.412, *n* = 2,442; Fig. 1c); meaning that many genes responsive to moderate stress also changed under RCP8.5, even when they did not pass our stringent DEG threshold. This asymmetry reveals threshold-like, non-linear shifts in gene regulation as climatic stress increases (Fig. 1d) and points to recruitment of additional molecular pathways under extreme conditions.

Functional enrichment analysis supports this interpretation. DEGs under RCP8.5 showed enrichment for Gene Ontology (GO) categories absent from RCP2.6 (Fig. 1e), while GO terms enriched under RCP2.6 were also overrepresented, though less strongly, under RCP8.5. The most significant RCP8.5-specific enrichments involved genes related to RNA modification, abscisic acid signaling, and negative regulation of phosphoprotein phosphatase activity. Downregulated genes were primarily linked to pathways for carbohydrate metabolism, lipid catabolism, and cell-wall organization, reflecting a shift in energy use and structural remodeling under extreme stress. In both RCP scenarios, upregulated DEGs were linked to gibberellin response, sterol metabolism, and ion transport, which may reflect subtle adjustments in growth and cellular homeostasis.

## Activation of unique transcriptional programs under RCP8.5

To explore the induction of unique regulatory pathways under the most severe climate scenario in more detail, we performed a weighted gene co-expression network analysis (WGCNA) across all the present (PC) and future climates (RCP2.6, RCP8.5). This approach grouped 19,873 genes into 28 co-expression modules (Fig. 2a) and enabled the identification of modules most associated with ambient CO_2_, ambient ozone, and stomatal ozone uptake (Fₒ_3_), as well as hub genes driving co-expression networks. The six modules exhibiting the strongest positive correlations with both CO_2_ and ozone (MEpink, MEgreenyellow, MEbrown) and strongest negative correlations (MEblue, MEdarkgreen, MEgrey60) contained genes that showed a strong uniform up- and down regulation across all samples under RCP8.5 (Fig. S1). In contrast, their expression patterns under RCP2.6 and PC were markedly weaker and more heterogeneous among samples (Fig. 2c, Supplementary Fig. S1). Furthermore, none of the 28 genes modules showed a significant association with stomatal ozone uptake (|r| < 0.2) (Fig. 2b), indicating that their gene expression was not driven by stomatal ozone flux. Subsequent functional enrichment analysis of these modules revealed a significant (q < 0.05) and exclusive enrichment of various RNA modification and processing functions in the MEbrown module (Fig. 2f), indicating that this module may function as a co-expressed RNA modification gene cluster. Notably, four of the five potential hub genes in this module encoded pentatricopeptide repeat-containing (PRP) proteins, Bhaga_6.g3124 (PRP At3g57430, chloroplastic), Bhaga_12.g2100 (PRP At3g12770-like), Bhaga_4.g2641 (PRP At4g21065-like), and Bhaga_5.g692 (PRP At3g14580, mitochondrial) (Fig. 2h). The strong uniform upregulation of this gene cluster suggests a major reorganization of post-transcriptional regulation under RCP8.5, patterns that were not visible in the other climate scenarios (Fig. 2e,f).

**Figure 2:**
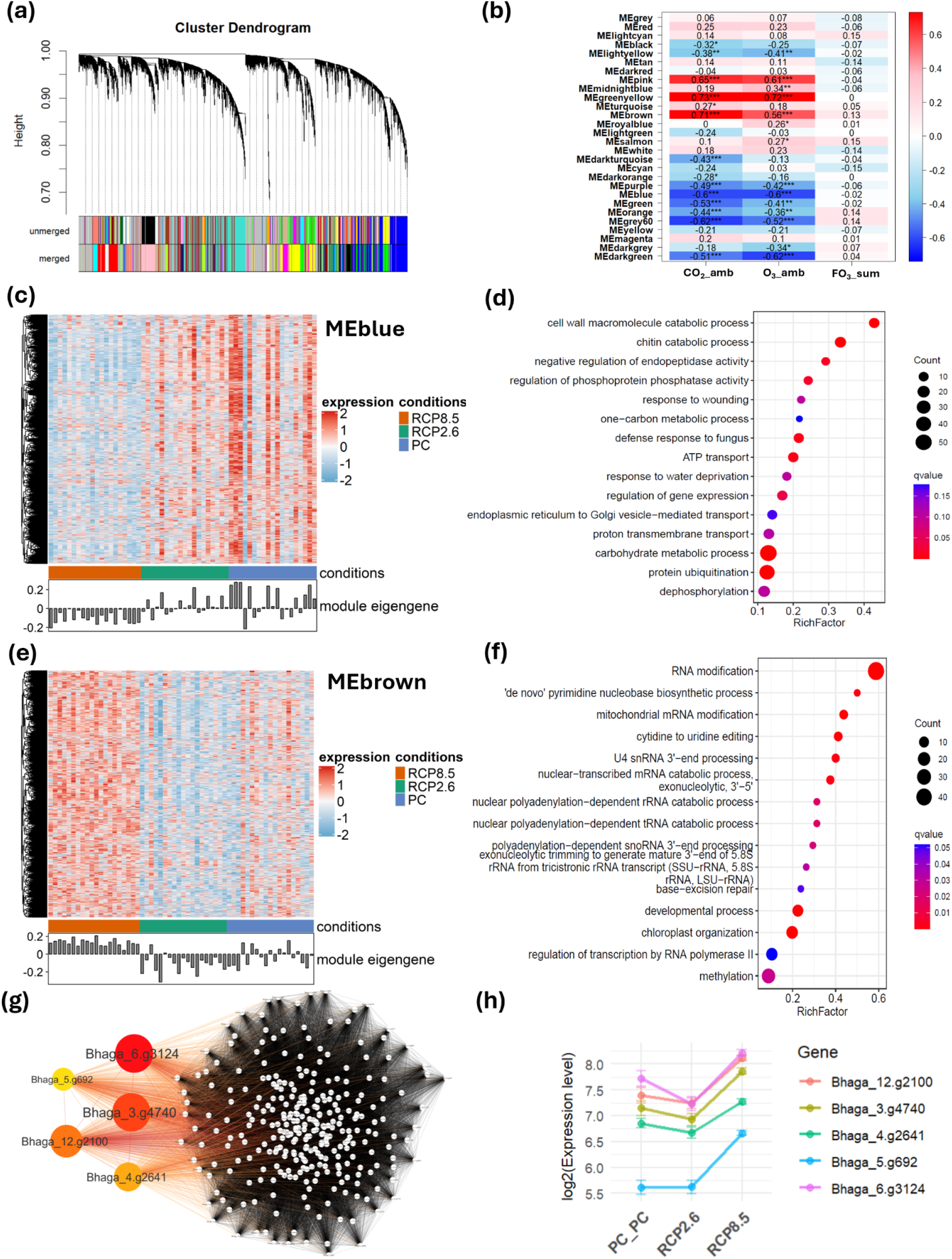
Activation of unique molecular pathways under PCP8.5. **(a)** Dendrogram of the hierarchical clustering of genes based on their TOM-based dissimilarity. Modules with module eigengene correlations (representative expression profiles) ≥ 0.75 were merged. **(b)** Correlation (Pearson) of the module eigengenes with ambient CO_2_ and O_3_ concentrations and sum of stomatal ozone uptake (FO_3_). **(c-h)** Gene expression heatmap, gene ontology analysis, and hub gene network of the blue (c-d) and the brown (e-h) module. **(c,e)** Heatmap visualization of module gene expression of each sample of the MEblue (c) and the MEbrown (e) module. **(d,f)** Gene ontology biological process (BP) overrepresentation analysis of the MEblue MEbrown (d) and the MEbrown module (f). RichFactor quantifies how enriched a specific term is compared to their respective background gene set. **(g)** Hub gene co-expression network based on co-expression similarity scores (TOM values), where node size represents the MCC rank of the hub genes and edge colors correspond to TOM weights. **(h)** Expression levels (log_2_) of identified hub genes of the brown module.

The MEblue module showed a strong negative correlation with both CO_2_ (r = –0.6) and ozone (r = –0.6) and was enriched in pathways encompassing carbohydrate metabolism and various cell wall macromolecule catabolic processes (Fig. 2c,d). While RCP2.6 and PC exhibited heterogeneous responses across samples, only RCP8.5 induced uniform downregulation, further highlighting a threshold-dependent transcriptional response that emerged exclusively under extreme climatic stress.

Collectively, these results show that while *F. sylvatica* exhibits a moderate transcriptional adjustment under milder RCP2.6 conditions, RCP8.5 triggers a distinct, more complex program, with RCP8.5-specific modules reflecting a threshold-like, non-linear integration of multiple environmental stressors at the transcriptome level.

## Discussion

Using a fully controlled ecotron experiment and high-resolution regionalized climate simulations, this study shows that *F. sylvatica* displays a pronounced non-linear transcriptional adjustment under RCP8.5, marked by a substantially intensified response and the activation of unique pathways and gene modules.

These emerging response patterns indicate that extreme future climates may elicit transcriptional dynamics that are not predictable from moderate-condition experiments. Moreover, elevated ozone alone induced only modest expression changes, and gene modules correlated weakly with stomatal ozone influx, suggesting that ozone was not the dominant driver. These results highlight that transcriptional responses arise from the combined action of multiple environmental factors, underscoring the need for realistic multifactorial experiments, as single-factor studies may underestimate future transcriptomic complexity.

The consistent and sharp downregulation of carbohydrate metabolism observed in all samples under RCP8.5 indicates that the elevated CO_2_ concentrations anticipated in this scenario may induce a saturation effect, where excess leaf carbohydrates trigger negative feedback due to limited sink capacity (Davey et al. 2006). Consistently, previous studies have reported downregulation of genes involved in carbohydrate biosynthesis and metabolism under elevated CO_2_, likely caused by excess carbon feedback (Huang et al. 2019; Kim et al. 2022).

Furthermore, under RCP8.5 conditions, *F. sylvatica* may strategically allocate resources toward rapid cell expansion and biomass accumulation in response to high-CO_2_ environments, as reflected in a strong downregulation of cell wall macromolecule catabolic processes and cell wall organization. Comparable responses have been documented in *Quercus petraea* and *Arabidopsis thaliana*, where CO_2_ enrichment altered cell wall composition, favoring cellulose over lignin, and increased cell wall thickness and mass (Mizokami et al. 2019; Koike et al. 2018; Teng et al. 2006).

The pronounced threshold-like activation of RNA modification pathways under RCP8.5 suggests that *F. sylvatica* rely on active transcriptome remodeling to sustain compensatory adaptive mechanisms under stronger climate forcing. Among the candidate regulators, pentatricopeptide repeat-containing proteins (PRPs) may play a key role in mediating this dosage-dependent response. PRPs are a large family of RNA-binding proteins essential for post-transcriptional organelle RNA metabolism (Laluk et al. 2011; Wang und Tan 2025), and some studies showed that PRPs exhibit sensitivity to biotic and abiotic stresses responses, likely mediated by reactive oxygen species (ROS) (Wang und Tan 2025). This may further indicate to an adaptive role of PRPs in acclimating *F. sylvatica* to RCP8.5 conditions, consistent with recent findings where PRP mutants exhibited significant ROS accumulation and enhanced sensitivity to salinity and ABA stress response in rice (Xiao et al. 2021) and *Arabidopsis (Zsigmond et al. 2008)*.

The strong activation of abscisic acid (ABA)-dependent signaling pathways may therefore represent an additional stress-response mechanism. As a central regulator of abiotic stress, ABA modulates stomatal closure to reduce water loss and enhances oxidative stress tolerance through antioxidant enzyme activation (Zhang et al. 2011; Tanotra et al. 2019). In line with this, the observed co-expression of negative regulation of Protein Phosphatase, suggests a reinforcement of ABA signaling pathways as part of *F. sylvatica*’s adaptive response under RCP8.5, as phosphatases are known to deactivate SNF1-Related Protein Kinase 2, the core drivers of ABA signaling (Umezawa et al. 2009).

Taken together, our study highlights the critical role of precisely controlled ecotron experiments, integrated with regionalized downscaled climate series, offering a unique approach to study the adaptive responses of *F. sylvatica* to the combined effects of future environmental conditions.

## Methods

### Plant material and growth

One year prior to the start of the experiment 5-10-year-old beech trees with average tree height of 110 ± 18 cm were taken from a natural regeneration forest site (Kranzberger Forest) together with a 20-liter soil monolith. After being planted into 25 L planters and acclimatized outdoors for one year with sufficient water supply, ten plants per climate scenario were transferred to precisely controlled ecotrons (Model EcoSystem Analyser, TUMmesa) engineered by regineering GmbH (Pollenfeld, Germany). Trees were then grown from April to September in the TUMmesa-Ecotron under four different climate regimes for two years (see section provision of simulation data for further details about the simulated climate scenarios). Trees were overwintered outside at the Greenhouses and Phytochambers Unit (GPU) of the TUM Plant Technology Center from October to March.

The ecotron system enables dynamic, minute-precise control of air temperature (ta), relative humidity (rh), light, CO_2_ and O_3_ as well as manipulation of soil temperature, soil moisture and nutrient supply. The LED system (Vossloh Schwabe, Urbach, Germany) provided a PPFD of >800 µmol m^−2^ s^−1^ within a near-natural UV-B to far-red spectrum. Plants were drip-irrigated daily to maintain 50 – 90% of the plant available soil water content, corresponding to 25.5 – 37.9% volumetric water content (VWC). VWC was monitored by a time domain reflectometer installed per planter. Air temperatures were controlled within a range of 4°C to 30°C. Highest possible rh was capped at 90%. A solution of 3.5 mM KH_4_NO_3_, 1.5 mM KNO_3_, 3 mM Ca(NO_3_)2,1.5 mM KH_2_PO_4_, 0.75 mM K_2_SO_4_, 0.9 mM MgSO_4_, 61 µM Fe-EDTA was applied weekly to each plant to ensure steady nutrient availability and the nutrient status of the plants were controlled in August of each growing season. For each sample ten leaves per tree were taken at the end of September of the second year (after 572 days) and were immediately frozen in liquid nitrogen and stored at −80°C. Each tree was sampled twice.

### RNA-Seq and data processing

RNA was extracted using the RNeasy Plant Mini Kit (Qiagen, Valencia, CA, USA) per the manufacturer’s instructions. The libraries were sequenced to 100 bp per read on a NextSeq500 platform (Illumina 1.9) by BGI. Adapter sequences and low-quality sequences were filtered out by SOAPnuke software developed by BGI. An average of 20,781,963 clean reads was obtained per sample (Table A. 1). The paired-end reads were then mapped to the reference genome of *Fagus sylvatica L. (Mishra et al. 2021; Mishra et al. 2018)* http://thines-lab.senckenberg.de/beechgenome/Bhaga_genome.fasta, accessed on Sep. 2024) using HISAT2 v2.2.1 (Kim et al. 2015) in order to generate splice-aware alignments. Alignment quality was assessed using samtools v.1.2.1 (Danecek et al. 2021) and RseQC v5.0.4 (Wang et al. 2012). Average alignment rate was 96.52% and gene body coverage showed uniform coverage across all samples and no 5’ or 3’ bias was detected. Gene expression counts were determined with featureCounts v.2.0.6 (Liao et al. 2014) and differentially expressed genes (DEGs) were identified by using DESeq2 2 v1.34.0 (Love et al. 2014). For identification of significant DEGs log2(fold change) values were used and an adjusted p-value of ≤ 0.05 was applied. Gene ontologies and functional annotation of the genes were obtained from the study by Mishra et al. (2021). Mishra et al. (2021) used a combination of protein queries against the NCBI non-redundant (NR) database (downloaded on June 24, 2020) using DIAMOND (v0.9.30) (Buchfink et al. 2015) to identify homologous sequences with known functions. Additionally, InterProScan (Jones et al. 2014) was employed to detect functional domains and sites. The final functional annotations of the genes were then assigned using Blast2GO (Götz et al. 2008). Gene ontology enrichment was performed by clusterProfiler v‘4.2.2’ in R (Yu et al. 2012) using a custom-made background gene set for the gene ontology terms. Enrichment was shown using the RichFactor, which measures the ratio of the selected input genes of the term against the total number of genes annotated to the term in the background gene set. For visualization purposes a variance stabilization transformation (vst) was applied to the expression counts to stabilize variance across different expression levels.

### Weighted Gene Co-Expression Analysis (WGCNA)

Count data was normalized using DESeq2 and variance stabilization transformation (vst) was applied. Furthermore, all genes with less than 15 counts in more than 75% of samples were discarded, resulting in a final set of 19,873 retained genes. WGCNA analysis was then conducted by using the WGCNA (v ‘1.73’) package in R (Langfelder und Horvath 2008). Based on the 19,873 genes from PC, RCP2.6, and RCP8.5 conditions, the softConnectivity function was utilized to calculate the mean connectivity, employing 5,000 randomly selected genes. A soft power threshold of 9 was chosen to ensure a relatively mean connectivity while still maintaining a high scale-free topology fit. Automatic network construction was performed using the blockwiseModules function utilizing a signed Topological Overlap Matrix (TOM). A merge cut height of 0.25 was applied to merge modules with eigengene correlation of ≥ 0.75. Module eigengenes of the modules are defined as the first principal component of the standardized expression data of all genes of the respective module (Langfelder und Horvath 2008), and thereby summarize the gene expression profiles within a module. Hierarchical clustering was performed using TOM-based dissimilarity (1-TOM). Module Eigengenes of the modules were correlated with the physical parameters and a Pearson correlation coefficient (|r| > 0.6) was applied to identify significantly associated modules. Module membership (MM) for each gene was determined by correlating gene expression with module eigengenes (MEs). Similarly, gene significance (GS) was assessed by correlating gene expression levels with physical parameters. For hub gene analysis, only significant genes from each significant module were selected, defined by a module membership (MM) and gene significance (GS) greater than 0.5. Edge weights were assigned based on topological overlap matrix (TOM) values, with a threshold of TOM > 0.05 applied for visualization. A continuous color gradient was used to represent the edge weights between nodes. Hub genes were identified using Maximal Clique Centrality (MCC) from the package Cytoscape package cytohubba (v0.1) (Chin et al. 2014), and the top 5 genes selected as hub genes based on their MCC rank.

### Climate simulation data

Three different realistic climate scenarios were generated for application in TUM*mesa*: a present climate (PC), representing a possible, average year of the 1987-2016 period at the reference climate station Spessart (Jossgrund-Lettgenbrunn, Germany, Code DEHE026, 497 m asl, 50°09′52.0″N 9°23′58.0″E) and two future climate scenarios. The future scenarios were developed in accordance with the representative concentration pathway (RCP) concept for the two contrasting scenarios RCP2.6 (van Vuuren et al. 2011) and RCP8.5 (Riahi et al. 2011)[BJ1] for the period 2071-2100. All three scenarios were calculated from the ReKliEs-DE (Heike Hübener et al. 2018) and EURO CORDEX (Jacob et al. 2014) climate model ensembles for a forested area in the German Spessart mountains that is characterized by managed beech and spruce stands. In order to generate realistic annual and diurnal courses in high temporal resolution, hourly resolved meteorological data were resampled from the historical record of the reference station in order to match statistical properties of the scenarios that were derived from the model ensembles. The procedure is described in detail by (Jákli et al. 2021b). Thus, climate series of air temperature (t_a_), relative humidity (rh) and global radiation (converted to PPFD according to (Grünhage und Haenel 1997)) were generated for each scenario. [O_3_] was derived by resampling for PC and adopted for the RCP scenarios following (Hayes et al. 2019): for RCP2.6, hourly values of [O_3_] concentration <40 ppb were increased by +5 ppb and [O_3_] >45 ppb were decreased by -5 ppb. For RCP8.5, [O_3_] <40 ppb was increased by +10 ppb and by +5 ppb for values between 40 and 50 ppb. [O_3_] ≥50 ppb was decreased by -5 ppb in the RCP8.5 scenario. The scenarios were complemented by a series of tropospheric CO_2_ concentrations that were adapted for PC, RCP2.6 and RCP8.5. First, a relative annual CO_2_ series was calculated by normalizing the average annual CO_2_ course recorded 1987-2016 at the climate station Schauinsland (1205 m asl, 47°54′49.7″N 7°54′27.9″E) on the respective average annual concentration of CO_2_. Representative time series for PC, RCP2.6, and RCP8.5 were generated by multiplying the relative hourly CO_2_ series with 375 ppm (30-years average at “Spessart” station), 421 ppm and 936 ppm (simulated for RCP2.6 and RCP8.5 in 2100, see (Meinshausen et al. 2011). To prevent pseudo-replication, trees and their corresponding climate scenarios were exchanged among the ecotron-chambers each year. The climate data was adapted for implementation in the TUM*mesa* climate chambers (Tab. 1) according to (Jákli et al. 2021b) and are available at (Jákli et al. 2021a). In order to isolate the O_3_-effect from compound climate effects, the present climate scenario PC was repeated with increased [O3]. Therefore, hourly [O_3_] of the PC were manipulated the same way as for RCP8.5.

### Ozone uptake (TransP Sensor)

Continuous measurement of stomatal ozone fluxes for beech was conducted using a TransP sensor (TransP, Ecomatik GmbH, Dachau, Germany), which measured the temperature difference between the underside of the beech leaf and the surrounding air. Using recently derived beech-specific calibration function, this TransP temperature signal was then used to determine the transpiration rate of water vapor (total leaf conductance). The total leaf conductance (mol m⁻² s⁻¹) was then converted to the ozone conductance by multiplying by 0.663, being the relative diffusivity of ozone compared to water vapor (Grünhage et al. 2012). Stomatal ozone flux in nmol m⁻² s⁻¹ was then determined by multiplying the ozone conductance with the external ozone concentrations. Hourly stomatal ozone flux measurements were recorded for every sample tree and time-integrated to obtain yearly total ozone uptake (FO3_sum, mmol m^-2^).

## Supplementary figures

**Figure S1:**
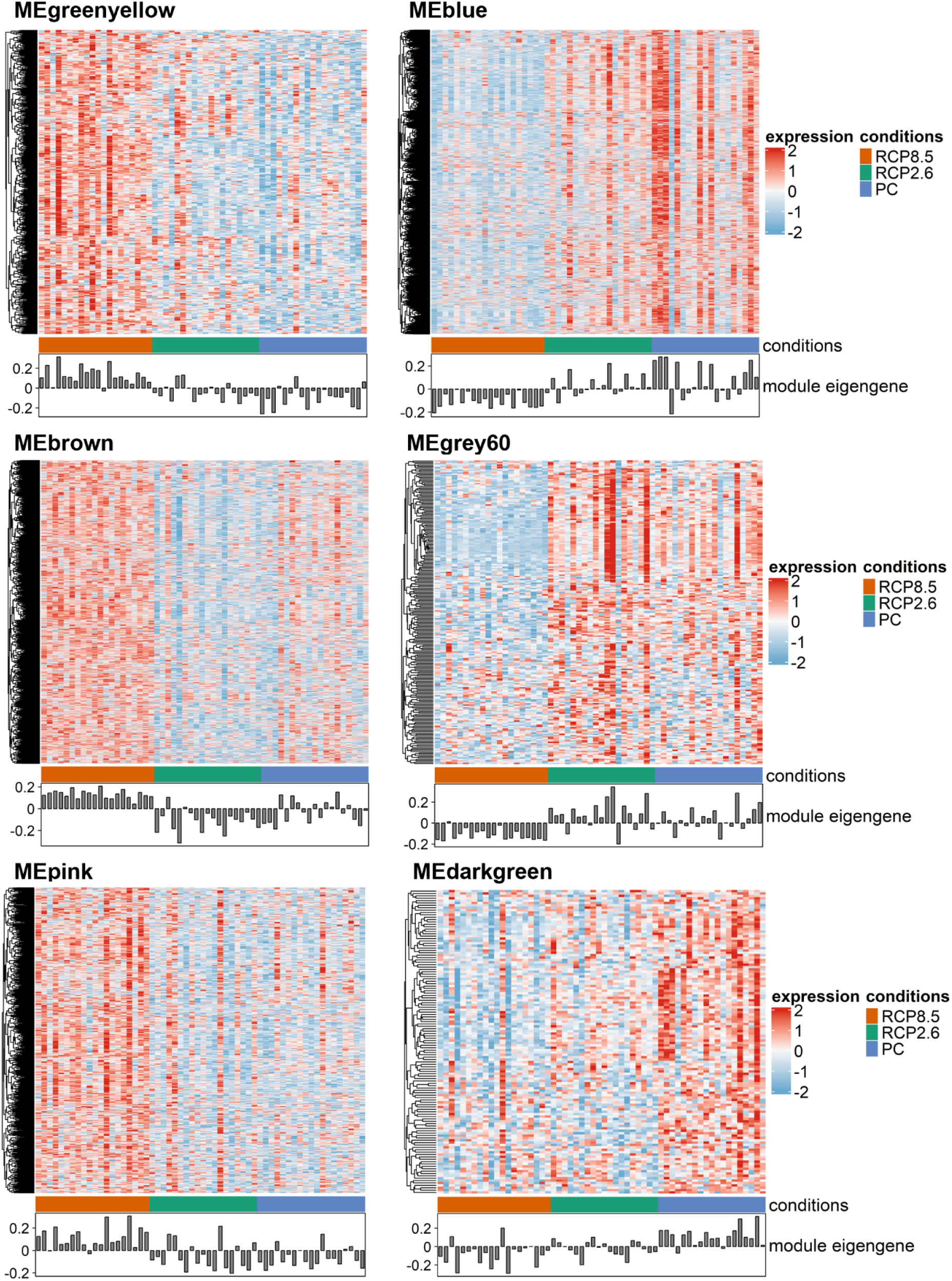
Heatmap visualization of the gene expressions (z-transformed) of each sample of the respective gene modules and their corresponding module eigengenes (representative gene expression profile).

**Figure S2:**
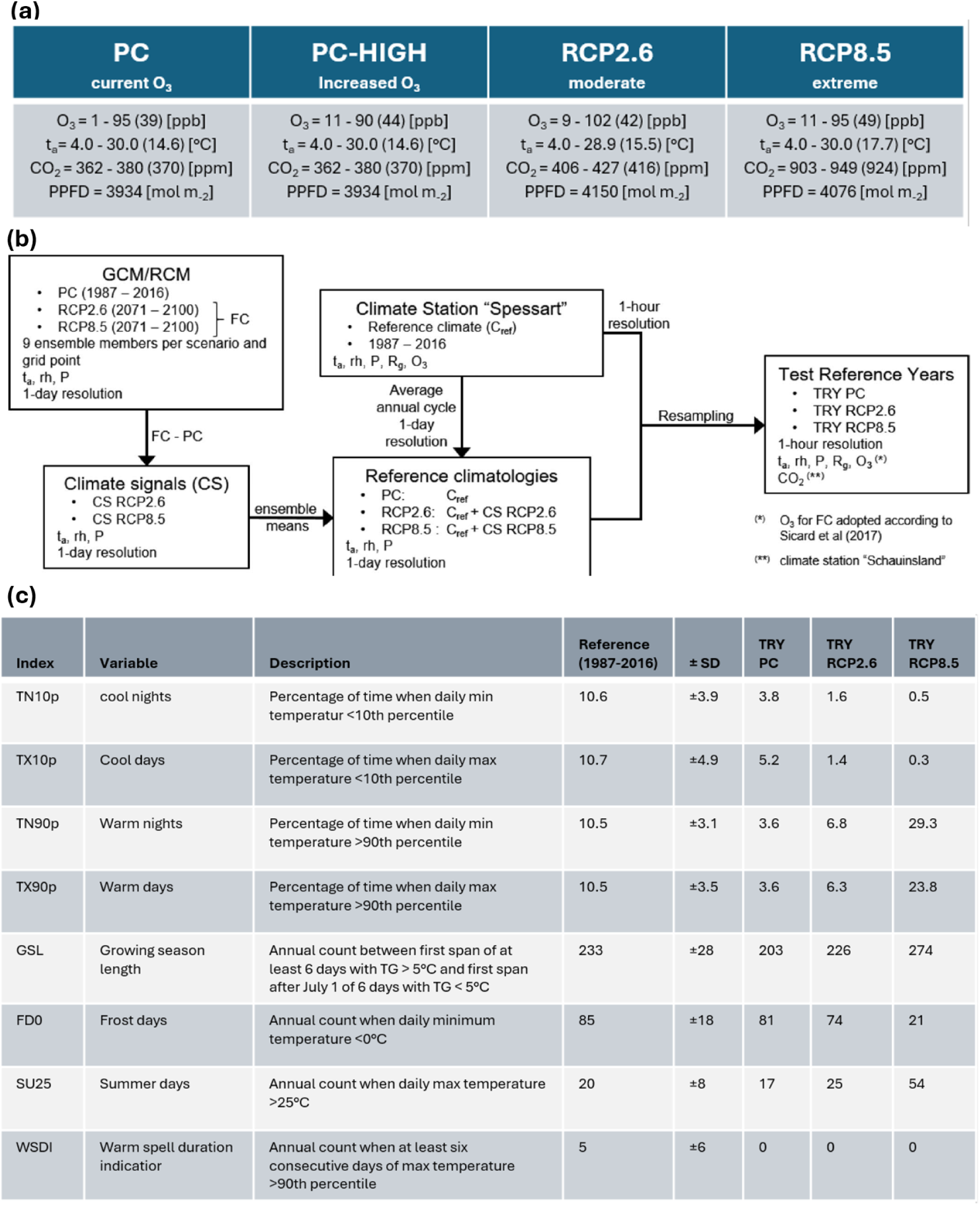
Climate time series workflow and key variables (a) range (mean) values of O_3,_ t_a_, CO_2_ and Photosynthetic Photon Flux Density (PPFD) of the simulated climate scenarios. (b) Schematic representation of the regionalized test reference year (TRY) generation. RCM ensembles were obtained and climate signals (CS) determined by subtracting FC values from PC for each RCM simulation (source: (Jákli et al. 2021b). Mean CS values were then added to the reference climate to create reference climatologies for the RCPs. Regional climate records were then resampled with the RCP reference climatologies to create regionalized TRYs. (c) Climate indices of the test reference years for PC, RCP2.6 and RCP8.5 recorded at the climate station „Spessart“.

## Bibliography

1. Ainsworth, Elizabeth A.; Long, Stephen P. (2005): What have we learned from 15 years of free-air CO2 enrichment (FACE)? A meta-analytic review of the responses of photosynthesis, canopy properties and plant production to rising CO2. In: The New phytologist 165 (2), S. 351–371. DOI: 10.1111/j.1469-8137.2004.01224.x.

2. Betz, G. A.; Gerstner, E.; Olbrich, M.; Winkler, J. B.; Langebartels, C.; Heller, W. et al. (2009): Effects of abiotic stress on gene transcription in European beech: ozone affects ethylene biosynthesis in saplings of Fagus sylvatica L. In: iForest 2 (3), S. 114–118. DOI: 10.3832/ifor0495-002.

3. Boeck, Hans J. de; Vicca, Sara; Roy, Jacques; Nijs, Ivan; Milcu, Alexandru; Kreyling, Juergen, et al. (2015): Global Change Experiments: Challenges and Opportunities. In: BioScience 65 (9), S. 922–931. DOI: 10.1093/biosci/biv099.

4. Buchfink, Benjamin; Xie, Chao; Huson, Daniel H. (2015): Fast and sensitive protein alignment using DIAMOND. In: Nature methods 12 (1), S. 59–60. DOI: 10.1038/nmeth.3176.

5. Cernusak, Lucas A.; Haverd, Vanessa; Brendel, Oliver; Le Thiec, Didier; Guehl, Jean-Marc; Cuntz, Matthias (2019): Robust Response of Terrestrial Plants to Rising CO2. In: Trends in plant science 24 (7), S. 578–586. DOI: 10.1016/j.tplants.2019.04.003.

6. Chin, Chia-Hao; Chen, Shu-Hwa; Wu, Hsin-Hung; Ho, Chin-Wen; Ko, Ming-Tat; Lin, Chung-Yen (2014): cytoHubba: identifying hub objects and sub-networks from complex interactome. In: BMC systems biology 8 Suppl 4 (Suppl 4), S11. DOI: 10.1186/1752-0509-8-S4-S11.

7. Clobert, Jean; Chanzy, André; Le Galliard, Jean-François; Chabbi, Abad; Greiveldinger, Lucile; Caquet, Thierry et al. (2018): How to Integrate Experimental Research Approaches in Ecological and Environmental Studies: AnaEE France as an Example. In: Front. Ecol. Evol. 6, S. 963. DOI: 10.3389/fevo.2018.00043.

8. Cooper, O. R.; Parrish, D. D.; Ziemke, J.; Balashov, N. V.; Cupeiro, M.; Galbally, I. E. et al. (2014): Global distribution and trends of tropospheric ozone: An observation-based review. In: Elementa: Science of the Anthropocene 2 (8), S. 1501. DOI: 10.12952/journal.elementa.000029.

9. Cuervo-Alarcon, Laura; Arend, Matthias; Müller, Markus; Sperisen, Christoph; Finkeldey, Reiner; Krutovsky, Konstantin V. (2021): A candidate gene association analysis identifies SNPs potentially involved in drought tolerance in European beech (Fagus sylvatica L.). In: Scientific reports 11 (1), S. 2386. DOI: 10.1038/s41598-021-81594-w.

10. Danecek, Petr; Bonfield, James K.; Liddle, Jennifer; Marshall, John; Ohan, Valeriu; Pollard, Martin O. et al. (2021): Twelve years of SAMtools and BCFtools. In: GigaScience 10 (2). DOI: 10.1093/gigascience/giab008.

11. Davey, P. A.; Olcer, H.; Zakhleniuk, O.; Bernacchi, C. J.; Calfapietra, C.; Long, S. P.; Raines, C. A. (2006): Can fast-growing plantation trees escape biochemical down-regulation of photosynthesis when grown throughout their complete production cycle in the open air under elevated carbon dioxide? In: Plant, cell & environment 29 (7), S. 1235–1244. DOI: 10.1111/j.1365-3040.2006.01503.x.

12. Garosi, Cesare; Ferrante, Roberta; Vettori, Cristina; Paffetti, Donatella (2022): Meta-Analysis as a Tool to Identify Candidate Genes Involved in the Fagus sylvatica L. Abiotic Stress Response. In: Forests 13 (2), S. 159. DOI: 10.3390/f13020159.

13. Götz, Stefan; García-Gómez, Juan Miguel; Terol, Javier; Williams, Tim D.; Nagaraj, Shivashankar H.; Nueda, María José et al. (2008): High-throughput functional annotation and data mining with the Blast2GO suite. In: Nucleic acids research 36 (10), S. 3420–3435. DOI: 10.1093/nar/gkn176.

14. Grünhage, L.; Haenel, H. D. (1997): PLATIN (plant-atmosphere interaction) I: A model of plant-atmosphere interaction for estimating absorbed doses of gaseous air pollutants. In: Environmental pollution (Barking, Essex : 1987) 98 (1), S. 37–50. DOI: 10.1016/S0269-7491(97)00114-0.

15. Grünhage, Ludger; Matyssek, Rainer; Häberle, Karl-Heinz; Wieser, Gerhard; Metzger, Ursula; Leuchner, Michael et al. (2012): Flux-based ozone risk assessment for adult beech forests. In: Trees 26 (6), S. 1713–1721. DOI: 10.1007/s00468-012-0716-5.

16. Günthardt-Goerg, Madeleine S.; Vollenweider, Pierre (2007): Linking stress with macroscopic and microscopic leaf response in trees: new diagnostic perspectives. In: Environmental pollution (Barking, Essex : 1987) 147 (3), S. 467–488. DOI: 10.1016/j.envpol.2006.08.033.

17. Gupta, Dharmendra K.; Palma, José M.; Corpas, Francisco J. (2015): Reactive Oxygen Species and Oxidative Damage in Plants Under Stress. Cham: Springer International Publishing.

18. Han, Qingmin; Kabeya, Daisuke; Hoch, Günter (2011): Leaf traits, shoot growth and seed production in mature Fagus sylvatica trees after 8 years of CO2 enrichment. In: Annals of botany 107 (8), S. 1405–1411. DOI: 10.1093/aob/mcr082.

19. Hanson, Paul J.; Walker, Anthony P. (2020): Advancing global change biology through experimental manipulations: Where have we been and where might we go? In: Global change biology 26 (1), S. 287–299. DOI: 10.1111/gcb.14894.

20. Hayes, Felicity; Mills, Gina; Alonso, Rocio; González-Fernández, Ignacio; Coyle, Mhairi; Grünhage, Ludger et al. (2019): A Site-Specific Analysis of the Implications of a Changing Ozone Profile and Climate for Stomatal Ozone Fluxes in Europe. In: Water Air Soil Pollut 230 (1). DOI: 10.1007/s11270-018-4057-x.

21. Heike Hübener; Katharina Bülow; Cornelia Fooken; Barbara Früh; Peter Hoffmann; Simona Höpp et al. (2018): ReKliEs-De Ergebnisbericht. FG Atmosphärische Prozesse.

22. Huang, Yulan; Fang, Rui; Li, Yansheng; Liu, Xiaobing; Wang, Guanghua; Yin, Kuide et al. (2019): Warming and elevated CO2 alter the transcriptomic response of maize (Zea mays L.) at the silking stage. In: Scientific reports 9 (1), S. 17948. DOI: 10.1038/s41598-019-54325-5.

23. IPCC (2023): IPCC, 2023: Climate Change 2023: Synthesis Report. Contribution of Working Groups I, II and III to the Sixth Assessment Report of the Intergovernmental Panel on Climate Change [Core Writing Team, H. Lee and J. Romero (eds.)]. IPCC, Geneva, Switzerland.

24. Unter Mitarbeit von Hoesung Lee, Katherine Calvin, Dipak Dasgupta, Gerhard Krinner, Aditi Mukherji, Peter W. Thorne et al.: Intergovernmental Panel on Climate Change (IPCC).

25. Jacob, Daniela; Petersen, Juliane; Eggert, Bastian; Alias, Antoinette; Christensen, Ole Bøssing; Bouwer, Laurens M. et al. (2014): EURO-CORDEX: new high-resolution climate change projections for European impact research. In: Reg Environ Change 14 (2), S. 563–578. DOI: 10.1007/s10113-013-0499-2.

26. Jákli, Bálint; Meier, Roman; Gelhardt, Ulrike; Bliss, Margaret; Grünhage, Ludger; Baumgarten, Manuela (2021a): Regionalized dynamic climate series for ecological climate impact research in modern controlled environment facilities.

27. Jákli, Bálint; Meier, Roman; Gelhardt, Ulrike; Bliss, Margaret; Grünhage, Ludger; Baumgarten, Manuela (2021b): Regionalized dynamic climate series for ecological climate impact research in modern controlled environment facilities. In: Ecology and evolution 11 (23), S. 17364–17380. DOI: 10.1002/ece3.8371.

28. Jones, Philip; Binns, David; Chang, Hsin-Yu; Fraser, Matthew; Li, Weizhong; McAnulla, Craig et al. (2014): InterProScan 5: genome-scale protein function classification. In: Bioinformatics (Oxford, England) 30 (9), S. 1236–1240. DOI: 10.1093/bioinformatics/btu031.

29. Karlsson, Per Erik; Klingberg, Jenny; Engardt, Magnuz; Andersson, Camilla; Langner, Joakim; Karlsson, Gunilla Pihl; Pleijel, Håkan (2017): Past, present and future concentrations of ground-level ozone and potential impacts on ecosystems and human health in northern Europe. In: The Science of the total environment 576, S. 22–35. DOI: 10.1016/j.scitotenv.2016.10.061.

30. Kim, Daehwan; Langmead, Ben; Salzberg, Steven L. (2015): HISAT: a fast spliced aligner with low memory requirements. In: Nature methods 12 (4), S. 357–360. DOI: 10.1038/nmeth.3317.

31. Kim, Tae-Lim; Lim, Hyemin; Chung, Hoyong; Veerappan, Karpagam; Oh, Changyoung (2022): Elevated CO2 Alters the Physiological and Transcriptome Responses of Pinus densiflora to Long-Term CO2 Exposure. In: Plants (Basel, Switzerland) 11 (24). DOI: 10.3390/plants11243530.

32. Koike, Takayoshi; Kitao, Mitsutoshi; Hikosaka, Kouki; Agathokleous, Evgenios; Watanabe, Yoko; Watanabe, Makoto et al. (2018): Photosynthetic and Photosynthesis-Related Responses of Japanese Native Trees to CO2: Results from Phytotrons, Open-Top Chambers, Natural CO2 Springs, and Free-Air CO2 Enrichment. In: William W. Adams III und Ichiro Terashima (Hg.): The Leaf: A Platform for Performing Photosynthesis, Bd. 44. Cham: Springer International Publishing (Advances in Photosynthesis and Respiration), S. 425–449.

33. Košvancová, M.; Urban, O.; Šprtová, M.; Hrstka, M.; Kalina, J.; Tomášková, I. et al. (2009): Photosynthetic induction in broadleaved Fagus sylvatica and coniferous Picea abies cultivated under ambient and elevated CO2 concentrations. In: Plant Science 177 (2), S. 123–130. DOI: 10.1016/j.plantsci.2009.04.005.

34. Laluk, Kristin; Abuqamar, Synan; Mengiste, Tesfaye (2011): The Arabidopsis mitochondria-localized pentatricopeptide repeat protein PGN functions in defense against necrotrophic fungi and abiotic stress tolerance. In: Plant physiology 156 (4), S. 2053–2068. DOI: 10.1104/pp.111.177501.

35. Langfelder, Peter; Horvath, Steve (2008): WGCNA: an R package for weighted correlation network analysis. In: BMC bioinformatics 9, S. 559. DOI: 10.1186/1471-2105-9-559.

36. Liao, Yang; Smyth, Gordon K.; Shi, Wei (2014): featureCounts: an efficient general purpose program for assigning sequence reads to genomic features. In: Bioinformatics (Oxford, England) 30 (7), S. 923–930. DOI: 10.1093/bioinformatics/btt656.

37. Love, Michael I.; Huber, Wolfgang; Anders, Simon (2014): Moderated estimation of fold change and dispersion for RNA-seq data with DESeq2. In: Genome biology 15 (12), S. 550. DOI: 10.1186/s13059-014-0550-8.

38. Martinez Del Castillo, Edurne; Zang, Christian S.; Buras, Allan; Hacket-Pain, Andrew; Esper, Jan; Serrano-Notivoli, Roberto, et al. (2022): Climate-change-driven growth decline of European beech forests. In: Communications biology 5 (1), S. 163. DOI: 10.1038/s42003-022-03107-3.

39. Meinshausen, Malte; Smith, S. J.; Calvin, K.; Daniel, J. S.; Kainuma, M. L. T.; Lamarque, J-F. et al. (2011): The RCP greenhouse gas concentrations and their extensions from 1765 to 2300. In: Climatic Change 109 (1-2), S. 213–241. DOI: 10.1007/s10584-011-0156-z.

40. Mishra, Bagdevi; Gupta, Deepak K.; Pfenninger, Markus; Hickler, Thomas; Langer, Ewald; Nam, Bora et al. (2018): A reference genome of the European beech (Fagus sylvatica L.). In: GigaScience 7 (6). DOI: 10.1093/gigascience/giy063.

41. Mishra, Bagdevi; Ulaszewski, Bartosz; Meger, Joanna; Aury, Jean-Marc; Bodénès, Catherine; Lesur-Kupin, Isabelle et al. (2021): A Chromosome-Level Genome Assembly of the European Beech (Fagus sylvatica) Reveals Anomalies for Organelle DNA Integration, Repeat Content and Distribution of SNPs. In: Frontiers in genetics 12, S. 691058. DOI: 10.3389/fgene.2021.691058.

42. Mizokami, Yusuke; Sugiura, Daisuke; Watanabe, Chihiro K. A.; Betsuyaku, Eriko; Inada, Noriko; Terashima, Ichiro (2019): Elevated CO2-induced changes in mesophyll conductance and anatomical traits in wild type and carbohydrate-metabolism mutants of Arabidopsis. In: Journal of experimental botany 70 (18), S. 4807–4818. DOI: 10.1093/jxb/erz208.

43. Müller, Markus; Seifert, Sarah; Lübbe, Torben; Leuschner, Christoph; Finkeldey, Reiner (2017): De novo transcriptome assembly and analysis of differential gene expression in response to drought in European beech. In: PloS one 12 (9), e0184167. DOI: 10.1371/journal.pone.0184167.

44. Ogle, Kiona; Liu, Yao; Vicca, Sara; Bahn, Michael (2021): A hierarchical, multivariate meta-analysis approach to synthesising global change experiments. In: The New phytologist 231 (6), S. 2382–2394. DOI: 10.1111/nph.17562.

45. Olbrich, M.; Betz, G.; Gerstner, E.; Langebartels, C.; Sandermann, H.; Ernst, D. (2005): Transcriptome analysis of ozone-responsive genes in leaves of European beech (Fagus sylvatica L.). In: Plant biology (Stuttgart, Germany) 7 (6), S. 670–676. DOI: 10.1055/s-2005-873001.

46. Olbrich, Maren; Gerstner, Elke; Bahnweg, Günther; Häberle, Karl-Heinz; Matyssek, Rainer; Welzl, Gerhard et al. (2010): Transcriptional signatures in leaves of adult European beech trees (Fagus sylvatica L.) in an experimentally enhanced free air ozone setting. In: Environmental pollution (Barking, Essex : 1987) 158 (4), S. 977–982. DOI: 10.1016/j.envpol.2009.08.001.

47. Paffetti, Donatella; Maren, Olbrich; Fladung, Matthias; Ernst, Dieter; Markussen, Thorsten; Forstreuter, Manfred et al. (2011): Effects of high levels of CO2 on gene expression in two different genotypes of Fagus sylvatica. In: BMC Proc 5 (S7). DOI: 10.1186/1753-6561-5-S7-P171.

48. Riahi, Keywan; Rao, Shilpa; Krey, Volker; Cho, Cheolhung; Chirkov, Vadim; Fischer, Guenther et al. (2011): RCP 8.5—A scenario of comparatively high greenhouse gas emissions. In: Climatic Change 109 (1-2), S. 33–57. DOI: 10.1007/s10584-011-0149-y.

49. Roy, Jacques; Rineau, François; Boeck, Hans J. de; Nijs, Ivan; Pütz, Thomas; Abiven, Samuel, et al. (2021): Ecotrons: Powerful and versatile ecosystem analysers for ecology, agronomy and environmental science. In: Global change biology 27 (7), S. 1387–1407. DOI: 10.1111/gcb.15471.

50. Seidl, Rupert; Thom, Dominik; Kautz, Markus; Martin-Benito, Dario; Peltoniemi, Mikko; Vacchiano, Giorgio et al. (2017): Forest disturbances under climate change. In: Nature climate change 7, S. 395–402. DOI: 10.1038/nclimate3303.

51. Tanotra, Suchita; Zhawar, Vikramjit Kaur; Sharma, Sucheta (2019): Regulation of Antioxidant Enzymes and Invertases by Hydrogen Peroxide and Nitric Oxide Under ABA and Water-Deficit Stress in Wheat. In: Agric Res 8 (4), S. 441–451. DOI: 10.1007/s40003-019-00399-6.

52. Teng, Nianjun; Wang, Jian; Chen, Tong; Wu, Xiaoqin; Wang, Yuhua; Lin, Jinxing (2006): Elevated CO2 induces physiological, biochemical and structural changes in leaves of Arabidopsis thaliana. In: The New phytologist 172 (1), S. 92–103. DOI: 10.1111/j.1469-8137.2006.01818.x.

53. Umezawa, Taishi; Sugiyama, Naoyuki; Mizoguchi, Masahide; Hayashi, Shimpei; Myouga, Fumiyoshi; Yamaguchi-Shinozaki, Kazuko et al. (2009): Type 2C protein phosphatases directly regulate abscisic acid-activated protein kinases in Arabidopsis. In: Proceedings of the National Academy of Sciences of the United States of America 106 (41), S. 17588–17593. DOI: 10.1073/pnas.0907095106.

54. UNEP (2023): Emissions Gap Report 2023: Broken Record – Temperatures hit new highs, yet world fails to cut emissions (again): United Nations Environment Programme.

55. van der Kooi, Casper J.; Reich, Martin; Löw, Markus; Kok, Luit J. de; Tausz, Michael (2016): Growth and yield stimulation under elevated CO 2 and drought: A meta-analysis on crops. In: Environmental and Experimental Botany 122 (1), S. 150–157. DOI: 10.1016/j.envexpbot.2015.10.004.

56. van Vuuren, Detlef P.; Edmonds, Jae; Kainuma, Mikiko; Riahi, Keywan; Thomson, Allison; Hibbard, Kathy et al. (2011): The representative concentration pathways: an overview. In: Climatic Change 109 (1-2), S. 5–31. DOI: 10.1007/s10584-011-0148-z.

57. Vanderkelen, Inne; Zscheischler, Jakob; Gudmundsson, Lukas; Keuler, Klaus; Rineau, Francois; Beenaerts, Natalie et al. (2020): A novel method for assessing climate change impacts in ecotron experiments. In: International journal of biometeorology 64 (10), S. 1709–1727. DOI: 10.1007/s00484-020-01951-8.

58. Vollenweider, P.; Ottiger, M.; Günthardt-Goerg, M. S. (2003): Validation of leaf ozone symptoms in natural vegetation using microscopical methods. In: Environmental pollution (Barking, Essex : 1987) 124 (1), S. 101–118. DOI: 10.1016/S0269-7491(02)00412-8.

59. Wang, Liguo; Wang, Shengqin; Li, Wei (2012): RSeQC: quality control of RNA-seq experiments. In: Bioinformatics (Oxford, England) 28 (16), S. 2184–2185. DOI: 10.1093/bioinformatics/bts356.

60. Wang, Yong; Tan, Bao-Cai (2025): Pentatricopeptide repeat proteins in plants: Cellular functions, action mechanisms, and potential applications. In: Plant communications 6 (2), S. 101203. DOI: 10.1016/j.xplc.2024.101203.

61. Xiao, Haijun; Liu, Zhongjie; Zou, Xue; Xu, Yanghong; Peng, Leilei; Hu, Jun; Lin, Honghui (2021): Silencing of rice PPR gene PPS1 exhibited enhanced sensibility to abiotic stress and remarkable accumulation of ROS. In: Journal of plant physiology 258-259, S. 153361. DOI: 10.1016/j.jplph.2020.153361.

62. Yu, Guangchuang; Wang, Li-Gen; Han, Yanyan; He, Qing-Yu (2012): clusterProfiler: an R package for comparing biological themes among gene clusters. In: Omics : a journal of integrative biology 16 (5), S. 284–287. DOI: 10.1089/omi.2011.0118.

63. Zhang, Aying; Zhang, Jun; Zhang, Jianhua; Ye, Nenghui; Zhang, Hong; Tan, Mingpu; Jiang, Mingyi (2011): Nitric oxide mediates brassinosteroid-induced ABA biosynthesis involved in oxidative stress tolerance in maize leaves. In: Plant & cell physiology 52 (1), S. 181–192. DOI: 10.1093/pcp/pcq187.

64. Zsigmond, Laura; Rigó, Gábor; Szarka, András; Székely, Gyöngyi; Otvös, Krisztina; Darula, Zsuzsanna et al. (2008): Arabidopsis PPR40 connects abiotic stress responses to mitochondrial electron transport. In: Plant physiology 146 (4), S. 1721–1737. DOI: 10.1104/pp.107.111260.

